# The Role of the Dorsolateral Prefrontal Cortex in Ego Dissolution and Emotional Arousal During the Psychedelic State

**DOI:** 10.1101/2024.12.09.627609

**Authors:** Clayton R. Coleman, Kenneth Shinozuka, Robert Tromm, Ottavia Dipasquale, Mendel Kaelen, Leor Roseman, Suresh Muthukumaraswamy, David J. Nutt, Lionel Barnett, Robin Carhart-Harris

## Abstract

Lysergic acid diethylamide (LSD) is a classic serotonergic psychedelic that induces a profoundly altered conscious state. In conjunction with psychological support, it is currently being explored as a treatment for generalized anxiety disorder and depression. The dorsolateral prefrontal cortex (DLPFC) is a brain region that is known to be involved in mood regulation and disorders; hypofunction in the left DLPFC is associated with depression. This study investigated the role of the DLPFC in the psycho-emotional effects of LSD with functional magnetic resonance imaging (fMRI) and magnetoencephalography (MEG) data of healthy human participants during the acute LSD experience. In the fMRI data, we measured the correlation between changes in resting-state functional connectivity (RSFC) of the DLPFC and post-scan subjective ratings of positive mood, emotional arousal, and ego dissolution. We found significant, positive correlations between ego dissolution and functional connectivity between the left & right DLPFC, thalamus, and a higher-order visual area, the fusiform face area (FFA). Additionally, emotional arousal was significantly associated with increased connectivity between the right DLPFC, intraparietal sulcus (IPS), and the salience network (SN). A confirmational “reverse” analysis, in which the outputs of the original RSFC analysis were used as input seeds, substantiated the role of the right DLPFC and the aforementioned regions in both ego dissolution and emotional arousal. Subsequently, we measured the effects of LSD on directed functional connectivity in MEG data that was source-localized to the input and output regions of both the original and reverse analyses. The Granger causality (GC) analysis revealed that LSD increased information flow between two nodes of the ‘ego dissolution network’, the thalamus and the DLPFC, in the theta band, substantiating the hypothesis that disruptions in thalamic gating underlie the experience of ego dissolution. Overall, this multimodal study elucidates a role for the DLPFC in LSD-induced states of consciousness and sheds more light on the brain basis of ego dissolution.

## 1. Introduction

Psychedelics have been utilized as therapeutic tools across the world for millennia(Carod-Artal, 2015). The revitalization of psychedelic research in the last decade has led to many new clinical trials, which have demonstrated that LSD and other psychedelic compounds may be promising treatments for psychiatric disorders. In conjunction with therapy, psilocybin is showing promise for treating end-of-life distress (Griffiths et al., 2016), smoking cessation (Johnson et al., 2017), major depressive disorder (MDD)(Carhart-Harris et al., 2021), and alcohol use disorder (Bogenschutz et al., 2022). Additionally, LSD has shown impressive results in clinical trials for generalized anxiety disorder (GAD) and MDD (Beutler et al., 2024; Holze et al., 2022).

Beyond psychedelics, there are other established interventions that may treat psychiatric conditions through potentially related mechanisms. An accelerated intermittent Theta Burst Stimulation (iTBS), a version of repeated Transcranial Magnetic Stimulation (rTMS), versus sham has shown an ∼90% response rate in limited clinical trials for the treatment of MDD (Cole et al., 2022). This intervention targets the left dorsolateral prefrontal cortex (lDLPFC) in a high-frequency, excitatory manner (Vida et al., 2023). A randomized sham-controlled study of rTMS excitation over the rDLPFC was found to effectively reduce symptoms of mania in conjunction with medication (Praharaj et al., 2009), and is now a third-line treatment for mania in Canada (Yatham et al., 2018). Moreover, low-frequency, inhibitory TMS over the *right* DLPFC (rDLPFC) is also effective for treating depression (Chen et al., 2013). These opposing effects of excitation and inhibition in the left and right DLPFC suggest a hemispheric lateralization of function within the DLPFC and its relevance to mood regulation and mood disorder. A recent meta-analysis showed that the left DLPFC plays a role in executive function while the right DLPFC has a stronger role in other aspects of cognitive processing and emotional responses (Lin & Feng, 2024).

Under LSD, ego dissolution strongly alters mood while also (acutely) resembling the mania-like symptoms of acute, first-episode psychosis (Preller & Vollenweider, 2018). Ego dissolution can either be euphoric or anxiety-inducing, and is characterized by experiences of interconnectedness, the blurring of the boundary between self and other, and a sense that ordinarily insignificant things have become profoundly meaningful (Friesen, 2022). Meanwhile, emotional arousal underpins the experience of mania and psychosis and is also a fundamental aspect of the psychedelic experience. Clinical populations with bipolar I disorder and schizophrenia typically experience difficulties with emotional regulation (Johnson et al., 2016), which can contribute to the experience of psychosis (Haralanova et al., 2012). In the psychedelic state, emotional arousal can manifest as euphoria, anxiety, or a loss of self-control, which can be examined through questionnaires such as the Altered States of Consciousness (ASC) scale (Dittrich, 1998).

In this study, we investigate whether the left and the right DLPFC play different roles in mediating the subjective experience of LSD. In a randomized, within-subjects, study of healthy volunteers, participants were administered LSD and placebo in two separate dosing sessions and asked to complete a post scan Visual Analogue Scale (VAS) including measures positive mood, ego dissolution, and emotional arousal. These were utilized to make correlations between these subjective ratings and resting-state functional connectivity (RSFC) of the left, right, and combined DLPFC, as captured with functional magnetic resonance imaging (fMRI). We predicted that changes in functional connectivity, of both the left and right DLPFC, induced by LSD would correlate with reported changes in ego dissolution, since psychedelics evoke a spectrum of effects that encompasses emotional arousal, including shifts in mood, and mania-like symptoms. Furthermore, we expected changes in lDLPFC seed-based FC under LSD would be associated with subject-reported changes in positive mood, based on the observation of changes in depressive symptoms resulting from TMS over the lDLPFC. Lastly, we expected alterations in rDLPFC under LSD to correlate with reported changes in emotional arousal, based on observed changes in arousal and mania-like symptoms after excitation of the rDLPFC in TMS.

## 2. Methods

### 2.1 Data Availability

LSD data was collected by the Centre for Psychedelic Research at Imperial College London (Carhart-Harris et al., 2020). The full dataset includes arterial spin labeling (ASL), functional magnetic resonance imaging (fMRI), and magnetoencephalography (MEG). We did not examine the ASL data in this analysis. A brief description of data collection and preprocessing procedures is given below, but the full methodology can be found in the primary paper (Carhart-Harris et al., 2016).

### 2.2 Participants

fMRI Blood Oxygen Level Dependent (BOLD) and MEG data were collected from 20 participants, of which 15 (four females; mean age, 30.5±8.0 y) were deemed suitable for analysis. One participant withdrew from the study due to excessive anxiety, while four others exhibited excessive head motion. For full inclusion criteria please refer to Carhart-Harris et al. (2016). Ethical approval was granted by the National Research Ethics Service committee London-West London. The research complied with the revised declaration of Helsinki (2000), the International Committee on Harmonization Good Clinical Practice guidelines, and National Health Service Research Governance Framework.

### 2.3 Study Design

Participants underwent two scanning sessions, one with placebo (PLCB) and one with LSD, separated by at least fourteen days. Participants were administered either PLCB, or 75 μg of LSD intravenously via a 10ml solution infused over two minutes. After a 60-minute acclimatization period, subjects were led to an MRI scanner for ASL and fMRI and then to magnetoencephalography (MEG) 165 minutes after drug administration. As stated above, the ASL data was not analyzed in this study. Both the fMRI and MEG recordings that were analyzed were eyes-closed resting-state scans. Eyes-open resting-state, eyes-closed music-listening, and video-watching recordings were also acquired, but were not included in this analysis due to a primary focus on eyes-closed rest (Carhart-Harris et al., 2016).

### 2.4 Subjective Reports

Directly after each scan, participants were administered a Visual Analogue Scale (VAS), which asks a series of questions about their subjective experience. The six domains of questions were the Intensity of the Experience, Simple Imagery, Complex Imagery, Positive Mood, Ego dissolution, and Emotional Arousal. The average (± S.D.) changes from PLCB to LSD were 10.4 ± 4.8 for Emotional Arousal, 6.7 ± 6.0 for Positive Mood, and 5.7 ± 7.2 for Ego Dissolution. At the end of each scan day, participants were also given the 11-dimensional Altered States of Consciousness (ASC) questionnaire, which asked participants to retrospectively rate their subjective experience at the peak of the compound’s effects.

### 2.5 fMRI

#### 2.5.1 fMRI Acquisition

MRI data was captured using a 3T GE HDx system. Two BOLD-weighted fMRI data sets were acquired via a gradient echo planar imaging sequence. Pulse sequence consisted of TR/TE = 2000/35ms, field-of-view = 220mm, 64 × 64 acquisition matrix, parallel acceleration factor = 2, 90° flip angle. Thirty-five oblique axial slices were captured in an interleaved manner, each being 3.4mm thick with zero slice gap (3.4mm isotropic voxels). Each of the two BOLD scans had a precise duration of 7:20 minutes. An additional 7:20 minute run was conducted between these two scans, but it was excluded from this study because of its added music component.

#### 2.5.2 fMRI Preprocessing

Preprocessing of the fMRI data was performed previously by Carhart-Harris et al. (2016) using a combination of four distinct yet complementary softwares: FMRIB Software Library (FSL)(Smith et al., 2004), AFNI (Cox, 1996), Freesurfer (Dale et al., 1999), and Advanced Normalization Tools (ANTS) (Avants et al., 2011). Participants were excluded if more than 15% of their volumes were scrubbed, with a scrubbing threshold of frame-wise head displacement (FWHD) = 0.5mm. Preprocessing stages included (1) removal of the first three volumes; (2) de-spiking; (3) slice time correction; (4) motion correction; (5) brain extraction; (6) rigid body registration to anatomical scans; (7) non-linear registration to 2mm MNI brain; (8) and scrubbing. Scrubbed volumes were replaced with the mean of the surrounding volumes.

Further pre-processing steps included (1) spatial smoothing; (2) band-pass filtering; (3) linear and quadratic detrending; (4) regression of nine nuisance regressors. Six nuisance regressors were motion-related and three were anatomically related. The three anatomical nuisance regressors calculated were ventricles (FreeSurfer, eroded in 2mm space), draining veins (FSL’s CSF minus FreeSurfer’s Ventricles, eroded in 1 mm space) and local white matter (WM) (FSL’s WM minus FreeSurfer’s subcortical gray matter (GM) structures, eroded in 2mm space). Regarding WM regression, AFNI’s 3dLocalstat was used to calculate the mean local WM time-series for each voxel, using a 25mm radius sphere centered on each voxel (Jo et al., 2010).

#### 2.5.3 fMRI Resting State Network Analysis

##### 2.5.3.1 Seed location

The seed-based analysis was performed with the use of FMRIB Software Library (FSL). Previous research of the DLPFC has utilized Brodmann Area 9/46, which does not account for recent developments in the understanding of the heterogeneity of this brain region (Cieslik et al., 2013; Jung et al., 2022). Thus, our study derived novel seed regions from rTMS studies that targeted the lDLPFC at (−42, 44, 30) in the Montreal Neurological Institute (MNI) coordinate system (Weigand et al., 2018). While this is an effective TMS stimulation point, it is not conducive for resting-state analysis because it is on the edge of the brain and is therefore prone to generating artifacts. To solve this problem, the lDLPFC seed was shifted by six units in each axis to (−36, 38, 24), thereby ensuring that an 8 mm sphere centered around it would not overlap with the edge of the brain. The rDLPFC seed was centered at the mirror location (36, 38, 24) in the right hemisphere.

##### 2.5.3.2 Functional Connectivity Analysis

Resting-state seed-to-voxel connectivity was measured between each brain voxel and the lDLPFC, rDLPFC, and combined DLPFC seeds. In the analyses of just the left or right DLPFC, the opposing seed’s activity was regressed out in the general linear model (GLM). We first analyzed the difference in connectivity between the PLCB and LSD conditions (Delta analysis), then examined correlations between the connectivity of each seed and the VAS ratings of Ego Dissolution, Emotional Arousal, and Positive Mood (Covariate analysis). All analyses were computed using FSL-Randomise (5000 permutations per test and contrast) (Winkler et al., 2014). Variance smoothing was employed at 6 mm for the between conditions *t-*test. Clusters were considered significant if *p*_FWE_ < 0.05, corrected for multiple comparisons using threshold-free cluster enhancement within trial (Smith & Nichols, 2009).

##### 2.5.3.3 Confirmational Analysis

The seeds in this study are novel; in particular, the individual and combined DLPFC seeds have traditionally not been used for resting-state FC analyses. Therefore, to ensure the robustness of our results, we performed a confirmational analysis in which we leveraged the absolute valued and binarized output clusters from the above analysis as the input seeds for new Delta and Covariate analyses. Our hypothesis was that the confirmational analysis would yield clusters containing the original DLPFC seeds. This would confirm that the subjective experiences of ego dissolution, emotional arousal, and positive mood are *specific* to the FC of the DLPFC.

### 2.6. MEG

#### 2.6.1 MEG Acquisition

Participants were recorded with a CTF 275-gradiometer MEG, though four of the sensors were turned off because of excessive sensor noise. Each scan lasted approximately seven minutes. MEG data was acquired at 600 Hz. There were two scans associated with the eyes-closed resting-state condition; we randomly selected one per subject. Simultaneous electrocorticography (ECG), vertical and horizontal electrooculography (EOG), and electromyography (EMG) recordings were acquired.

#### 2.6.2 MEG Preprocessing and Source Reconstruction

Preprocessing of the MEG data in this study was similar, but not identical, to the pipeline described in (Carhart-Harris et al., 2016). All preprocessing was performed in FieldTrip (Oostenveld et al., 2011). Data was high-pass filtered at 1 Hz with a 2^nd^-order Butterworth filter, downsampled initially to 400 Hz, and segmented into 2-second epochs. Line noise at 50 and 100 Hz was removed with a Discrete Fourier Transform filter. Outlier epochs and channels were manually deleted by visual inspection. An automatic algorithm was used to remove muscle artifacts from right-hemisphere temporal channels at the edge of the MEG dewar, which are most likely to be contaminated by such artifacts (Muthukumaraswamy et al., 2015). In particular, such channels with high activity (*z* > 6) and frequency between 105 and 145 Hz were suppressed. Independent component analysis (ICA) using the logistic infomax algorithm was applied to detect ECG and EOG artifacts (Bell & Sejnowski, 1995). Components were manually removed by visual inspection. Finally, data was downsampled again to 200 Hz to reduce the size of the data. The key differences between our preprocessing pipeline and that of Carhart-Harris et al. (2016) are the lack of *automatic* ICA artifact removal and the downsampling to 200 Hz.

Source reconstruction followed a procedure similar to the one described in (Mediano et al., 2024). The fMRI analysis yielded four networks: sets of regions that correlated with ratings of ego dissolution and emotional arousal, in two different “directions” (“forward,” or original, and “reverse,” or confirmational). For each network, a template consisting of the centroids of the constituent regions was inversely warped to each subject’s native-space anatomical MRI. A head model for each participant was constructed using the single-shell method. Based on the corresponding leadfield model, a linearly constrained minimum variance (LCMV) beamformer was applied to the inversely-warped template, with the regularization parameter set to 5% of the average of the diagonal elements of the sensor covariance matrix (Van Veen et al., 1997).

#### 2.6.3 Granger Causality (GC) Analysis

Directed functional connectivity was estimated between the constituent regions in each network with Granger causality. Broadly speaking, GC describes the extent to which the past activity of brain region *x* predicts the future of brain region *y* above and beyond the past of *y* (Granger, 1963, 1969).

Here, GC estimation was based on linear autoregressive (AR) modelling of the data (Barnett & Seth, 2014; Geweke, 1984), and computed from the AR model parameters by a state-space method (Barnett & Seth, 2015; Solo, 2016) [An AR model may be readily transformed into an equivalent innovations-form state-space model (Hannan & Deistler, 2012)] which devolves to the solution of a Discrete-time Algebraic Riccati Equation (DARE). Model orders (number of lags) for the AR models were selected using the Hannan-Quin information criterion (Hannan & Quinn, 1979) and model parameters obtained via Ordinary Least Squares (OLS). Pairwise-conditional GCs between sources (i.e., GCs between pairs of sources conditioned on the remaining sources in the network) were computed in the time domain, and in the frequency domain at a spectral resolution of 1024; frequency-domain GCs were then integrated across the frequency ranges 0-4 Hz, 4-8 Hz, 8-13 Hz, 13-30 Hz, 30-48 Hz, and 48-100 Hz to obtain band-limited GC (Barnett & Seth, 2011) in the delta, theta, alpha, beta, low-gamma, and high-gamma bands, respectively. As a sanity check, we verified that band-limited Granger causality values summed to the corresponding time-domain GC. All of these analyses were conducted with the Multivariate Granger Causality toolbox, version 2.0 (MVGC-2) (Barnett & Seth, 2014).

Time domain and band-limited GCs were computed for each of the four networks, in both experimental conditions (placebo and LSD). In the time domain, within-condition statistical significance against a null hypothesis of vanishing GC was evaluated using an F-test (Barnett & Seth, 2011). For band-limited GCs, statistical inference was evaluated by permutation testing (500 epoch-wise permutations), since an analytical sampling distribution for conditional band-limited GC remains unknown. An asymptotic sampling distribution in the *unconditional* case has recently been obtained by Gutknecht & Barnett (2023). Subject-level *p*-values were aggregated at the group level by Fisher’s method. Between conditions, Wilcoxon signed-rank tests were used to compare time-domain and band-limited GCs [delta and high gamma were excluded from this analysis due to the segmentation of the data into 2-second epochs, which is the length of one slow delta cycle, and the presence of artifacts in high gamma MEG data (Muthukumaraswamy, 2013)]. In all significance tests, the Benjamini-Hochberg False Discovery Rate (FDR) was applied to correct for multiple comparisons; in the band-limited case, a second level of FDR correction was applied to adjust for multiple comparisons across the four frequency bands.

## 3. Results

### 3.1 Delta Analysis (fMRI)

In the “Delta” analysis, we first measured the effects of LSD on seed-to-voxel resting-state functional connectivity (RSFC) between the combined, bilateral DLPFC seed and the rest of the brain, irrespective of the correlation with subjective ratings. This analysis revealed that LSD increased connectivity between the seed and the two major hubs of the Default Mode Network (DMN), the medial prefrontal cortex and the posterior cingulate cortex. It also revealed a reduction in RSFC between this seed and cortical regions including the left angular gyrus, right supramarginal gyrus, left precuneus, and a region in the left DLPFC. As the DLPFC is part of the frontoparietal network (FPN), this result substantiates the hypothesis that psychedelics increase between-network connectivity between the FPN and the DMN (Carhart-Harris, Muthukumaraswamy, et al., 2016; Girn et al., 2023; Müller et al., 2018; Roseman et al., 2014).

We also measured the RSFC of the left and right DLPFC on their own. LSD significantly reduced the RSFC between the lDLPFC and the rest of the brain, especially in the sensorimotor cortex (**Supplementary Figure 1**). (Note that global signal regression was not performed on the data.) LSD did not have a significant effect on the RSFC of the rDLPFC.

### 3.2 Covariate Analysis (fMRI)

We first examined correlations between the connectivity of the combined DLPFC (both left and right hemisphere, **Figure 1**) and subjective ratings of ego dissolution, emotional arousal, and positive mood. We found significant correlations between ego dissolution and two clusters with peaks at the left thalamus and the right fusiform gyrus (rFG) (**Figure 3a**). Given the novel nature of our original seeds, we sought to test the robustness of this correlation. Therefore, we conducted a confirmational analysis, in which the thalamus and rFG were the seed regions. If the analysis returned a cluster within the DLPFC, it would verify the specificity of the DLPFC in ego dissolution. Indeed, this confirmational analysis revealed two main clusters, which contained not only the rDLPFC but also the rIFG (**Figure 3b**). A smaller cluster, not visible in the figure, encompasses the left DLPFC, but outside the established seed.

**Figure 1:**
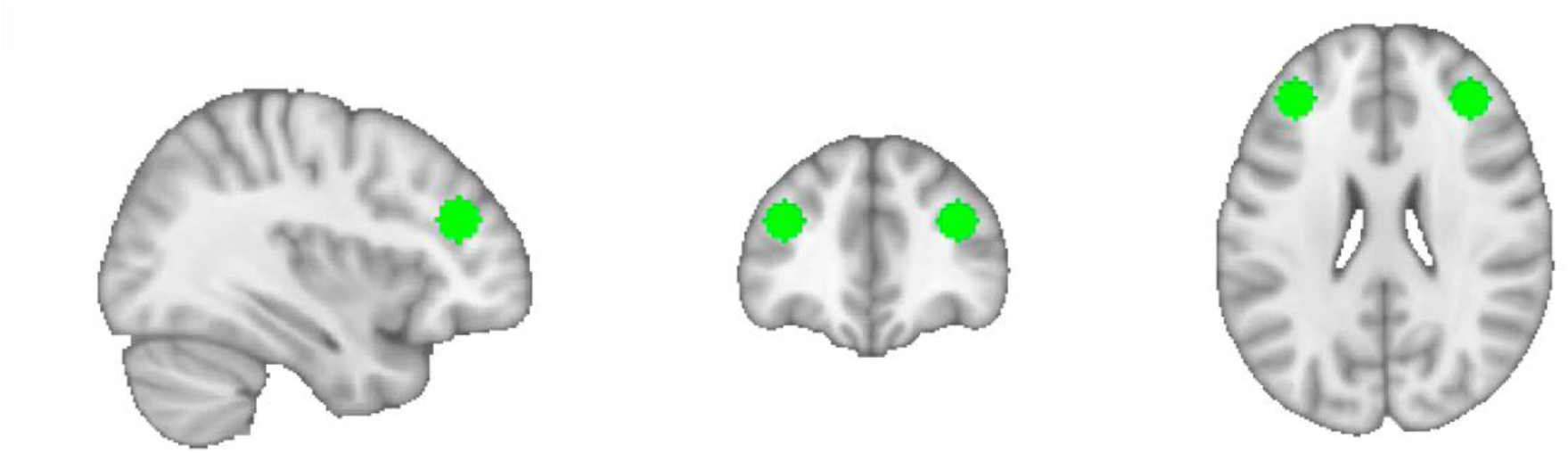
The left and right seeds of the dorsolateral prefrontal cortex (DLPFC) that were used in this study. Resting-state functional connectivity (RSFC) analyses were conducted on the combined, bilateral DLPFC, as well as on the left and right DLPFC separately.

**Figure 2:**
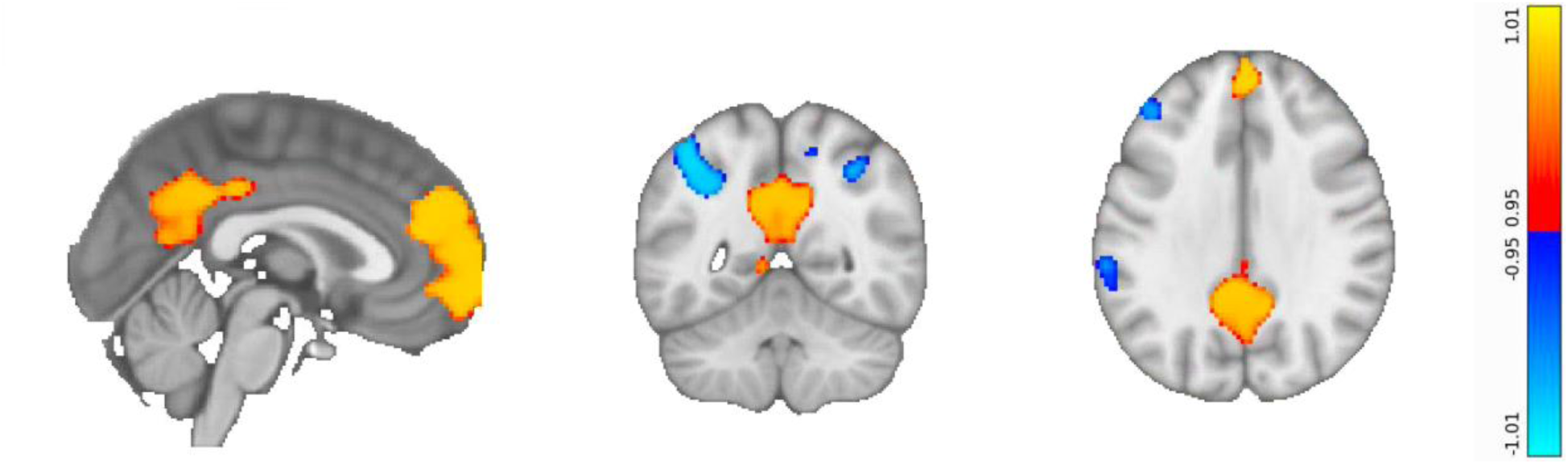
Delta analysis reveals regions of the brain that become more or less functionally connected to the combined DLPFC on LSD. Regions that increase in RSFC on LSD are shown in red/yellow; they include key hubs of the default mode network, i.e., the medial prefrontal cortex and posterior cingulate cortex. Blue areas denote regions that decrease in RSFC on LSD; they include the left angular gyrus, right supramarginal gyrus, left precuneus, and part of the left DLPFC.

**Figure 3:**
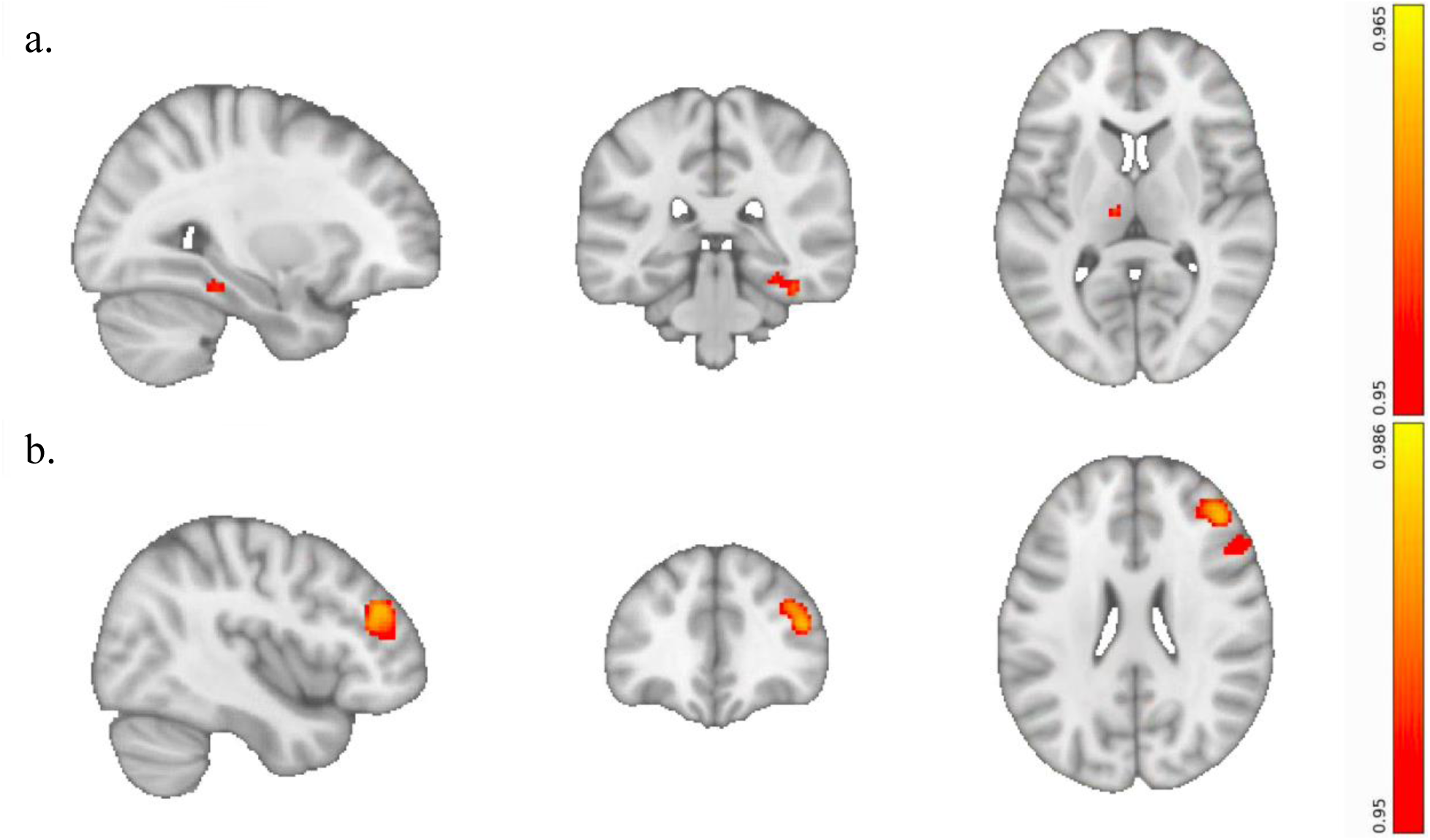
Ego dissolution ratings significantly correlate with the RSFC of the bilateral DLPFC. a) The connectivity between the bilateral DLPFC and the right fusiform gyrus (FG) & left thalamus is positively associated with ego dissolution i.e., greater coupling between the DLPFC and these regions, the greater the ego-dissolution. b) In the “reverse,” or confirmational, analysis, we made the outputs of the previous analysis the inputs for a new RSFC analysis. That is, we measured the correlation between ego dissolution and the RSFC of the right FG and left thalamus. We expected the reverse analysis to output the bilateral DLPFC. The analysis yielded clusters in the rDLPFC and right inferior frontal gyrus (rIFG) (detailed results in the Supplementary Materials).

Emotional arousal significantly correlated with connectivity between the rDLPFC and the IPS (**Figure 4a**). The confirmational analysis reproduced a portion of the rDLPFC while also revealing new regions, namely the right and left anterior insula (rAI, lAI), dorsal anterior cingulate cortex (dACC), and middle temporal gyrus (MTG), that correlated with emotional arousal (**Figure 4b**). The rAI, lAI, and dACC are main nodes of the Salience Network.

**Figure 4:**
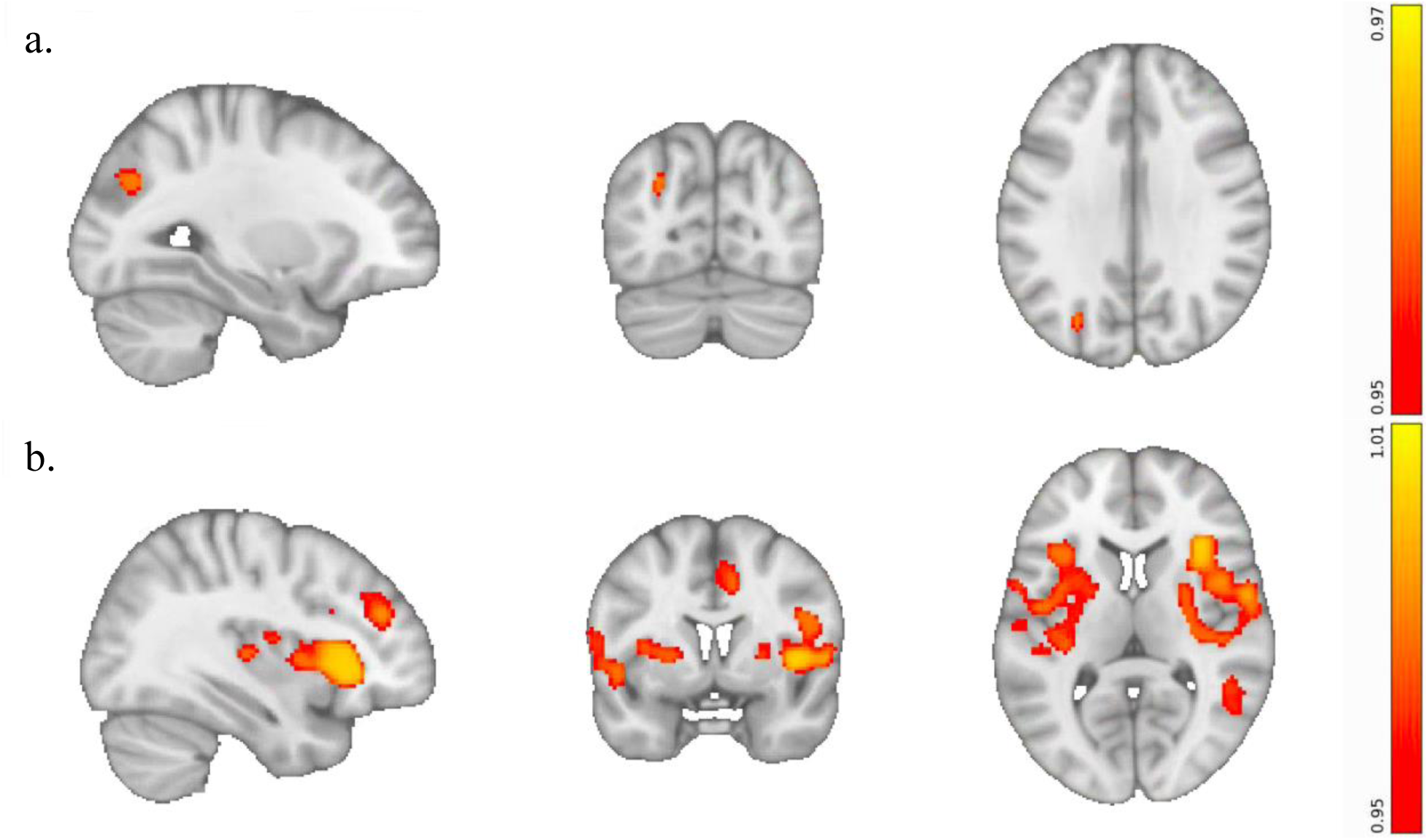
The RSFC of the right but not left DLPFC is significantly correlated with ratings of emotional arousal. a) RSFC between the rDLPFC and the left Intraparietal Sulcus (IPS) is positively associated with emotional arousal. b) Confirmational analysis, in which the rDLPFC and IPS were used as input seeds, yields clusters in the rDLPFC, middle temporal gyrus, and key nodes in the salience network, namely the left and right anterior insula and dorsal anterior cingulate cortex.

**Figure 5:**
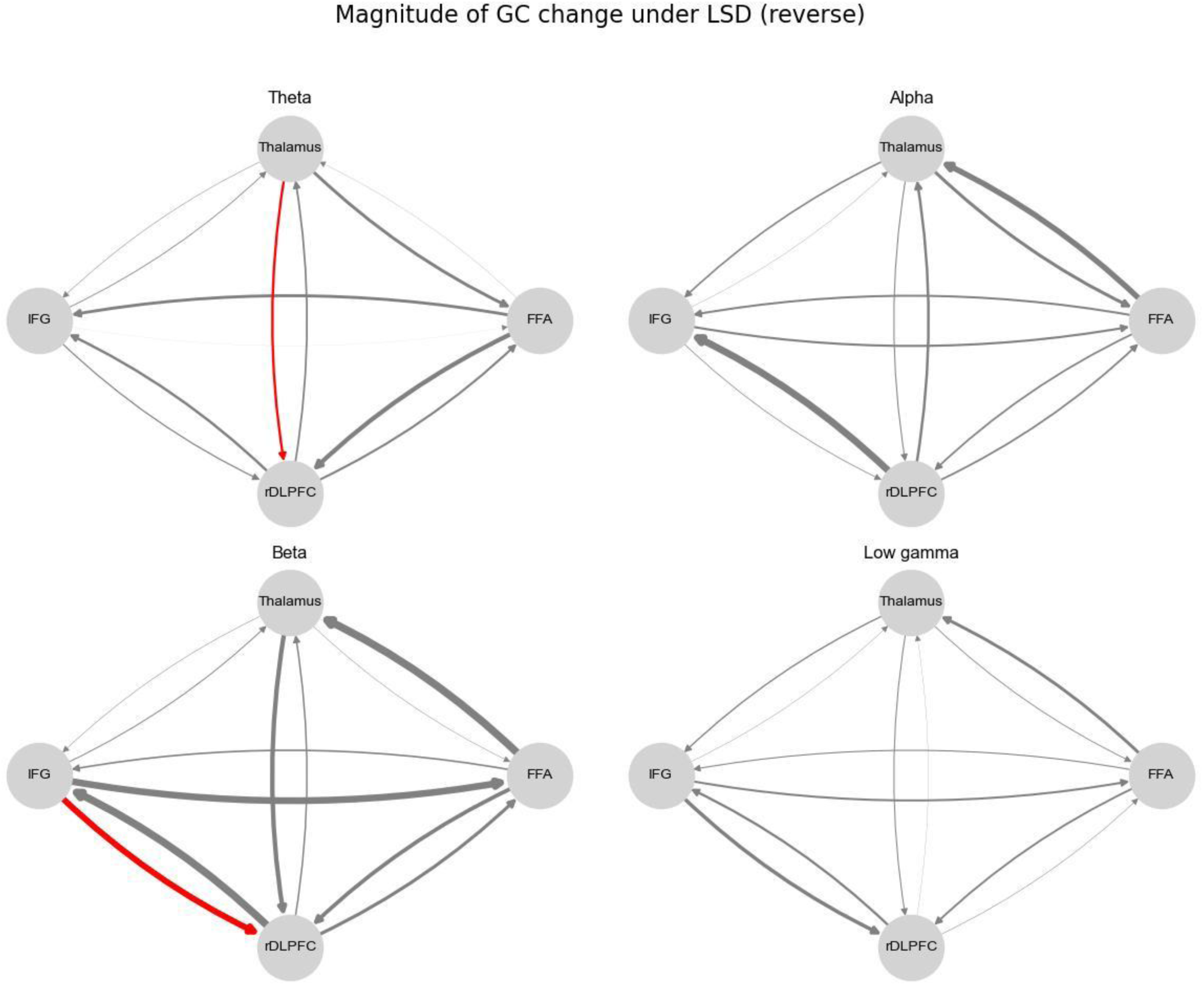
Magnitude of changes in GC between conditions for forward and reverse analyses of clusters correlating with ego dissolution. Significant increase (red) in GC from thalamus to rDLPFC in the theta band (p < 0.05, FDR corrected) and from IFG to rDLPFC in the beta band (p < 0.05, FDR corrected) for ego reverse analysis.

We did not find any significant correlations between positive mood and connectivity of the left, right, or combined DLPFC.

### 3.3 GC Analysis (MEG)

Within each network, we measured Granger Causality (GC), an estimate of directed functional connectivity, between each pair of constituent regions, conditioned on all other regions in the network. We computed both time-domain (broadband) and frequency-domain GC in each network.

Within each condition (placebo and LSD), all pairwise-conditional time-domain GC values were significant for all four of the networks (*p* < 0.002 in all cases). All frequency-domain GC values were significant in all four of the frequency bands – theta, alpha, beta, and low gamma – in all of the networks. Between placebo and LSD, there were no significant differences in time-domain GC for any pair of regions in any of the networks. However, frequency-domain GC did exhibit significant differences between conditions. In particular, LSD significantly increased theta-band GC from the thalamus to the rDLPFC and beta-band GC from the IFG to the rDLPFC in the ego reverse network (*p* = 0.0366). There were no significant between-condition differences in frequency-domain GC for the ego forward, emotional forward or emotional reverse network. Our results suggest that LSD specifically alters GC in a portion of the rDLPFC, i.e., the one in the ego reverse network, rather than the entire rDLPFC seed that was used in the ego forward network.

## 4. Discussion

In this study, we examined the effect of LSD on the functional connectivity of the left, right, and combined DLPFC, as measured with fMRI in healthy human participants. Then, we explored associations between subjective ratings of the LSD experience and the FC of each DLPFC seed. In the fMRI analysis, we find that LSD overall increases FC between the combined DLPFC and the DMN. Furthermore, the covariate of ego dissolution is positively correlated with an increased FC between the combined DLPFC and rFG and thalamus, as well as between rFG and thalamus and the rDLFPC and rIFG. On the other hand, emotional arousal is positively correlated with an increased FC between the rDLPFC and IPS, as well as between the IPS and the rDLPFC and nodes within the SN. Our complementary analysis of GC in the MEG dataset revealed that LSD significantly increases bottom-up connectivity from the left thalamus to the rDLPFC in the theta-band, as well as beta-band connectivity from the rIFG to the rDLFPC.

### 4.1 Lateralization of Function in the DLPFC

Our study was motivated by the evidence that TMS therapy and psychedelic-assisted therapy report similar therapeutic outcomes via two different treatment modalities. If both methodologies can reduce subjective ratings of depression, they may do so through the engagement of similar mechanisms. TMS for depression has evolved from a multitude of targets to a specific coordinate in the lDLPFC (Cash et al., 2021). We utilized a generalized coordinate and its mirror in the right hemisphere as the seeds for our RSFC analyses (Weigand et al., 2018). This data could then be correlated with subjective reports of the acute LSD experience, namely VAS ratings of ego dissolution, emotional arousal, and positive mood. Through our forward and reverse analyses, we were able to confirm the importance of the rDLPFC in the subjective experiences of emotional arousal and ego dissolution, two intertwined but separable experiences.

These results contribute to a relatively new understanding of the lateralization of function between the left and right DLPFC. As stated in the Introduction, this lateralization is exhibited in the different therapeutic effects associated with stimulating the left versus the right DLPFC with TMS. In particular, while stimulating the left DLPFC can reduce the severity of depression symptoms, stimulating the right DLPFC can inhibit mania. Our findings regarding the neural correlates of ego dissolution and emotional arousal are consistent with the lateralization of function in the DLPFC as revealed by TMS. We also found that the rDLPFC is particularly implicated in bottom-up shifts in network balance.

We propose that the function of primarily the right, and not the left, DLPFC shapes the experience of emotional arousal under psychedelics, which explains our finding that emotional arousal significantly correlates with the RSFC of only the rDLPFC. We contend that emotional arousal on LSD corresponds to a state phenomenologically similar to mania, particularly with regard to the variety of emotional valence. Emotional arousal captures both positive and negative changes in emotion on psychedelics; it is an unvalenced measure of subjective experience. This generalizable phenomenon is best understood as an umbrella term that can encompass multiple dimensions, which can be associated with positive or negative feelings. Secondly, one could construe increases in emotional arousal under LSD as consistent with an increase in perceived ‘salience’ (Friesen, 2022; Wießner et al., 2023), thus invoking the “aberrant salience” model of mania (Sass & Pienkos, 2013) and psychosis more generally (Kapur, 2003).

On the other hand, we propose that ego dissolution recruits the functions of *both* the left and right DLPFC, hence ego dissolution is correlated with the RSFC of both hemispheres. Indeed, ego dissolution can be highly euphoric yet also resemble a state resembling manic, first-episode psychosis, where ‘self’ abnormalities are known to be common (Moe, 2016; Sass & Pienkos, 2013). Ego dissolution predicts increases in positive emotional reactivity to psychedelic use (Orłowski et al., 2022) and can also predict improved well-being. Note also that a “mystical” loss of boundaries between the self and the other can also characterize manic psychosis. People experiencing this state have described a “breach in the barriers of individuality” leading to a sense of spiritual communion with the universe (John Custance, 1952; Landis, 1964; Parnas et al., 2005; Sass & Pienkos, 2013, 2013). In some cases, this state can trigger beliefs of, for instance, telepathy, or other such so-called “narcissistic delusions” such as “delusions of reference”. The phenomenology of this state is very similar to that of ego dissolution on psychedelics, which can also sometimes give rise to claimed telepathic-like experiences (Johnstad, 2020).

### 4.2 The DLPFC and Emotional Arousal

Our fMRI RSFC forward analysis found significant correlations between emotional arousal and connectivity between the rDLPFC and left IPS. The IPS is thought to be involved in regulating emotional arousal during threats. For instance, an fMRI and MEG study found global brain connectivity increases in the left IPS when analyzing threat detection (Balderston et al., 2017). Low frequency rTMS over either left or right IPS during threat of shock have been shown to reduce arousal (Balderston et al., 2020). These findings have led to a theory of connectivity in which the IPS mediates hyperarousal in anxiety disorders through DLPFC regulation (Brown et al., 2023).

The reverse analysis, in which the IPS cluster was used as the seed, returned a cluster in the rDLPFC. While this cluster was in a different part of the rDLPFC and smaller than the original seed, it nevertheless confirmed that the rDLPFC is specifically involved in mediating emotional arousal on LSD. The outputs of the reverse analysis also contain clusters in the bilateral AI, dACC, and MTG. The lAI, rAI, and dACC are well-known hubs of the Salience Network (SN), which processes the emotional significance or salience of information. Previous work has shown that the SN acts as an intermediary between the central executive network (CEN) and the DMN. Some recent work also indicated a potential over-involvement of the SN in depression (Lynch et al., 2024). Hubs in these three networks show differences in information flow and dominance of signaling contingent on the task(Menon & Uddin, 2010; Molnar-Szakacs & Uddin, 2022). Aberrations in network transitions are theorized to be at the root of psychological illness (Menon, 2011). Furthermore, as explained in *Section 4.1*, experiences of aberrant salience are a key aspect of emotional arousal on LSD; emotional sensitivity to the environment is heightened because everything appears to be extremely significant. Future work on this network may take into account the DMN for GC analysis or examine individual dyads within the network for other subjective categories like anxiety.

Our MEG analysis of these forward and reverse networks displayed statistically significant connections between all pairs of regions, across all frequency bands combined, under both placebo and LSD, which would suggest that regions form a genuine network in the brain. However, there were no statistically significant changes from placebo to LSD in emotional arousal in the frequency or time domain. This suggests that changes in directed information flow result not from broadband signals, but specific oscillatory patterns, which has been proposed by previous research (Clarke-Williams et al., 2024; Vinck et al., 2023). Further research should investigate the role of this circuit in self-other relations and ego dissolution under altered states of consciousness.

### 4.3 The DLPFC and Ego Dissolution

The experience of ego dissolution, or “the loss of a sense of oneself,” is an essential aspect of the classical psychedelic experience (Millière, 2017). Furthermore, there is a relationship between ego dissolution and statistically significant reductions in depression scores within adults, even in the long term (Carhart-Harris, Kaelen, et al., 2016; Roseman et al., 2018; Weiss et al., 2024). Hence, reliable biomarkers and mechanisms of the psychedelic experience of ego dissolution are worth further investigation. By identifying the network of regions that mediate the experience, we may be able to stimulate them with TMS, either independently of or in conjunction with a psychedelic treatment, to augment ego dissolution and thereby improve or better predict the therapeutic outcomes (Copa et al., 2024)

The psychedelic experience of ego dissolution has been correlated with brain activity in a wealth of previous research. A previous RSFC study, which was conducted on the same LSD data examined in this paper, revealed that global connectivity changes in the left and right angular gyrus and left and right insula correlated with ego dissolution (Tagliazucchi et al., 2016). Ego dissolution has mainly been attributed to connectivity changes within the DMN (Carhart-Harris, Muthukumaraswamy, et al., 2016; Lebedev et al., 2015) but also reduced interhemispheric connectivity within the salience network (Lebedev et al., 2015). This ‘disintegration’ of the high-level networks has been conceived as a ‘relaxation’ of top-down inhibitory control, which, according to the so-called ‘RElaxed Beliefs Under pSychedelics’ (REBUS) model, demonstrates a “flattening” of the functional hierarchy in the brain and corresponding ‘relaxed beliefs’ or assumptions about the self and environment. In a variety of mental illnesses, including depression, beliefs or assumptions can become (pathologically) over-weighted; thus, their relaxation under (and potentially after) psychedelic experience may explain their potential therapeutic utility (Moe, 2016; Sass & Pienkos, 2013). Previous work has made significant strides toward establishing changes in functional cortical hierarchy under psychedelics (Girn et al., 2022; Luppi et al., 2021; Shinozuka et al., 2024).

This study correlated ego dissolution with a network of brain regions, rather than a single region or the whole brain. We began by selecting a novel seed that comprised the combined left and right DLPFC. We identified a significant correlation between ego dissolution and changes in RSFC between the combined DLPFC and the thalamus & FFA. A confirmational, or reverse analysis, in which the FFA & thalamus became the input seeds, yielded two main outputs: a smaller rDLPFC cluster and the rIFG.

These analyses, which were performed in fMRI, capture undirected connectivity. We then measured directed connectivity in an MEG dataset that was acquired on the same set of participants. In particular, we evaluated GC on the source-reconstructed timeseries of the regions in the forward and reverse networks. While many methods exist for capturing directed connectivity, GC has strong advantages in identifying time-delayed interactions, is robust to noisy data, and is flexible in working with the frequency domain. While we did not directly correlate GC with the ego dissolution VAS scales, this analysis nevertheless revealed the effect of LSD on the direction of information flow between the identified clusters across different frequency bands: theta, alpha, beta, and low gamma. While the effects of LSD on directed FC across the whole brain have been previously documented (Barnett et al., 2020), our analysis reveals changes in information flow specific to areas correlated with ego dissolution. Our findings are inconsistent with previous literature, which found that LSD exclusively decreased GC (Barnett et al., 2020). However, our estimates of GC were conditional on all other regions in the network, whereas previous analyses were either unconditional or conditional on timeseries that were averaged across multiple regions.

The reverse analysis revealed two statistically significant increases in GC: from the thalamus to the rDLPFC in the theta band and from the IFG to the rDLPFC in the beta band. The centroid of the thalamus seed is specifically located within the medial dorsal (MD) thalamus, which is structurally and functionally connected to the DLPFC. In fact, the primary source of input to the parvocellular MD is the DLPFC (Byne et al., 2009; Mitchell & Chakraborty, 2013; Pergola et al., 2015). Increased bottom-up information flow from the thalamus to higher-order regions of the brain, such as the prefrontal cortex, has been proposed as a critical component of the neural mechanisms underlying the psychedelic experience (Avram et al., 2021; Onofrj et al., 2023; Vollenweider & Geyer, 2001). The thalamus plays a role in gating sensory information to the cortex, screening out irrelevant stimuli (McCormick & Bal, 1994).

Aberrant connectivity from the thalamus to the cortex can lead to a “sensory overload,” inundating the cortex with excessive interoceptive and exteroceptive information (Vollenweider & Geyer, 2001). Altered connectivity between the thalamus and the DLPFC is a robust biomarker of schizophrenia and psychosis, conditions in which patients assign too much meaning to irrelevant stimuli (Anticevic et al., 2014; Pergola et al., 2018; Steullet, 2020; Welsh et al., 2010; Woodward et al., 2012). Similarly, some studies have shown that psychedelics exclusively increase functional connectivity between the thalamus and many regions of cortex (Bedford et al., 2023; Müller et al., 2017; Tagliazucchi et al., 2016), though other studies have found some decreases in thalamocortical connectivity(Avram et al., 2022; Gaddis et al., 2022; Preller et al., 2018, 2020). Psilocybin specifically alters activity in a region of the thalamus that overlaps most with the MD nucleus, which is consistent with our findings (Gaddis et al., 2022). A recent theory proposes that theta-band spiking in the thalamus shifts thalamocortical coupling to a dysrhythmic state that underlies altered states of consciousness, which is also aligned with our observations of elevated theta-band thalamocortical connectivity on LSD (Onofrj et al., 2023).

The DLPFC is generally involved in determining whether information is pertinent enough to enter and stay in working memory (Altamura et al., 2010; Friedman & Robbins, 2022; Petrides, 2000; Rassi et al., 2023; Rosero Pahi et al., 2020). Crucially, the ability of the PFC to maintain and assign relevance to neural representations is mediated by the MD thalamus (Bolkan et al., 2017; Marton et al., 2018; Mitchell & Chakraborty, 2013; Parnaudeau et al., 2015; Rikhye et al., 2018; Schmitt et al., 2017) Rodent studies have demonstrated that MD neurons stabilize context-relevant representations in the PFC, thereby enabling them to be maintained within working memory, while suppressing context-irrelevant representations (Rikhye et al., 2018).

Thus, enhanced directed connectivity from the MD thalamus may cause information from typically deemed irrelevant contexts to interfere with representations in the PFC, leading to a “hyper-flexible” state in which the brain is less constrained by the demands of the present cognitive context. We speculate that, during self-reflection, heightened connectivity between the MD thalamus and the cortex may amplify neural representations that do not ordinarily pertain to the self. These amplifications could induce instability across this network including thalamic dependent internal sensory perception within the rIFG (Dobrushina et al., 2021) and distort functionality in any one of the other regions in the ego dissolution network, which have all been correlated with self-identification (Herwig et al., 2012; Kaplan et al., 2008; Ma & Han, 2012). This broadening of relevance could explain the experience of the “loss of the sense of self” found in ego dissolution; on psychedelics the neural mechanisms of the self “over-represent” or assign too much relevance to external stimuli, yielding a sense of vast interconnectedness with the environment.

Furthermore, increased beta-band connectivity from the rIFG to the rDLPFC, as we observed, could be involved in integrating, or attempting to integrate, experiences of ego dissolution. In particular, the rIFG and the rDLPFC are both part of a network that integrates discomfirmatory evidence, i.e., present stimuli that violate a prior belief (Ehlis et al., 2024; Lavigne et al., 2015). In other words, when prior held beliefs about the self are challenged by conflicting evidence, the rIFG and the rDLPFC work together to integrate this information and thereby maintain or update the belief. This could be consistent not only with the rIFG’s established role in integrating internal representations (Adolfi et al., 2017; Dobrushina et al., 2021), but also REBUS’ hypothesis that psychedelics revise prior beliefs by altering top-down inhibition from higher-order regions like the rIFG and the rDLPFC.

The observed effects of LSD on directed connectivity are similar to the neural signatures of schizophrenia, which relates to a longstanding, but contentious, view that psychedelics are “psychotomimetics” or models of psychosis (Nichols & Walter, 2021; Umbricht et al., 2003; Vollenweider et al., 1998). Schizophrenia and mania are associated with a number of structural deficits in the DLPFC, including cortical thinning, reduced gray matter volume, and lower fractional anisotropy of outgoing white matter tracts (Abé et al., 2023; Ellison-Wright & Bullmore, 2009; van Haren et al., 2011). Furthermore, as stated above, aberrations in connectivity between the MD thalamus and the PFC are commonly observed in patients with schizophrenia, perhaps explaining observations that task-irrelevant representations interfere more with cognition in these populations (Wagner et al., 2013).

While some of the changes in brain activity and phenomenology on LSD are also found in individuals with schizophrenia, it is important to note that schizophrenia is a chronic condition, whereas the LSD trip is a transient experience. Also, disruptions to self-awareness in schizophrenia differ significantly from those of LSD. For instance, patients over-attribute agency to themselves and overestimate their causal power over the world, whereas people who take LSD tend not to do so (Hur et al., 2014). Furthermore, individuals affected by schizophrenia often exhibit long-standing emotional withdrawal and cognitive blunting, which are markedly different from the emotional arousal and cognitive flexibility experienced on LSD (Doss et al., 2021; Martínez et al., 2021; Oorschot et al., 2013).

In summary, we propose that amplified connectivity from the thalamus to the DLPFC may cause representations of the outside world to interfere with self-related cognition, thereby giving rise to the experience of ego dissolution. The same neural mechanisms may underpin delusional beliefs about the self in patients afflicted with schizophrenia and psychosis.

### 4.4 Limitations

Because the data is resting-state, it is difficult to relate different patterns of brain activity to specific psychological behaviors or processes that are modulated via LSD. To provide stronger evidence, tasks must be deployed to examine changes in self-awareness in response to working memory demands, while acknowledging the risk of generalized deficit confounds in this regard.

We did not correlate the directed connectivity of the MEG data to ratings of ego dissolution because these ratings were not available for all of the included participants. However, because the undirected connectivity between the same regions did correlate with ego dissolution in the fMRI dataset, we are confident that the directed connectivity plays some role in shaping the experience of ego dissolution. That being said, whole-brain MEG analyses may have uncovered a broader set of brain regions that correlate with ego dissolution and emotional arousal.

Finally, accurate source reconstruction of MEG data to subcortical regions like the thalamus is generally challenging, especially in resting-state data (Krishnaswamy et al., 2017). fMRI analysis of subcortical activity may be more reliable, so dynamic causal modeling on the fMRI data, which can be used to estimate directed connectivity, may be worth exploring to assess the robustness of the MEG GC findings.

## 5. Conclusion

This study set out to show that changes in functional connectivity with the dorsolateral prefrontal cortex correspond to the subjective effects of the psychedelic state. In the fMRI analysis, ratings of emotional arousal correlated with connectivity between the salience network & IPS, and the right, but not left, DLPFC, implying a lateralization of function in the DLPFC. On the other hand, ego dissolution was associated with increased connectivity between the combined left & right DLPFC and clusters in the thalamus & the fusiform gyrus. A GC analysis of these networks elucidated the directionality of information flow between the constituent regions of each network. LSD elevated theta-band GC from the thalamus to the rDLPFC, which exemplifies an increase in bottom-up information flow and the flattening of the hierarchy of the brain on psychedelics.

Our study of both undirected and directed connectivity of the DLPFC on LSD has unveiled new methods and opportunities for research in psychedelic science. Firstly, distinguishing the lDLPFC and rDLPFC could improve understanding of not only psychedelics but also psychiatric disorders. In particular, analyzing the two hemispheres separately may reveal the lateralization of the DLPFC’s function, whereas measuring their activity together could elucidate the extent to which the two hemispheres function in unison. Finally, studies informed by the structural heterogeneity of the DLPFC and its receptor makeup could further elucidate this region’s role in the psychedelic experience and psychiatric illnesses. Secondly, this is one of the first studies to analyze the effects of psychedelics with both fMRI and MEG data. Future studies on psychedelics could leverage this multimodal approach, which takes advantage of the high spatial resolution of fMRI and high temporal resolution of MEG (Hall et al., 2014). Finally, the confirmational analysis played an essential role in our understanding of these networks, enabling us to ensure the validity of our novel seeds and substantiating the lateralization of function in the DLPFC.

Our findings provide novel insights into the neural mechanisms of ego dissolution and other altered states of consciousness on psychedelics, which may be similar to the neural pathways that are affected in psychosis and schizophrenia. Future research could determine whether the same neural mechanisms play a role in the therapeutic effects of psychedelics for disorders such as depression. Perhaps the reason ego dissolution is correlated with a reduction in depression is that, for a moment in time, the self-perpetuating definition of ‘who we believe we are’ stops long enough for us to envision broader possibilities and recreate ourselves.

## Data and Code Availability

Code is available upon request by contacting the corresponding author at. Processed fMRI data is open-source and available at https://openneuro.org/datasets/ds003059/versions/1.0.0. MEG data may be available upon request by contacting Leor Roseman at.

## Author Contributions

C.R.C. conceptualized the study, conducted all fMRI analyses, created Figures 1-3, and wrote the manuscript.

K.S. preprocessed and source-reconstructed the MEG data, conducted most of the Granger causality analyses on the MEG data, and wrote the manuscript together with C.R.C.

R.T. advised on analyses, created Figure 4, and edited the manuscript.

L.R., S.M., D.J.N., and R.C-H conceptualized the fMRI and MEG experiments and collected the data.

L.B. provided essential guidance on the Granger causality analysis, including help with writing scripts and performing the within-condition significance tests.

M.H and O.D. served as master’s thesis advisors for C.R.C. and oversaw the fMRI analyses.

## Declaration of Competing Interests

The authors have no competing interests to declare.

## Acknowledgments

C.R.C would like to express sincere gratitude to his thesis advisors Matt Howard and Ottavia Dipasquale and the staff at King’s College London IoPPN—especially Fernando Zelaya, Owen O’Daly, and Eamonn Walsh—for their invaluable support. Special thanks to Mitul Mehta for his thesis review and for suggesting the conformational analysis.

C.R.C. is funded by the Usona Institute. K.S. is funded by the Clarendon Fund, a Department of Psychiatry studentship, the John Henry Jones Scholarship from Balliol College, the Research Council U.K., and the Usona Institute. R.T. is funded by the Usona Institute. R.C-H is funded by a Ralph Metzner endowment. L.B. is supported by European Research Council Advanced Investigator Grant CONSCIOUS; grant number 101019254

## Notes

https://openneuro.org/datasets/ds003059/versions/1.0.0

### Competing Interest Statement

The authors have declared no competing interest.

