## Supplementary Figures for "The Role of the Dorsolateral Prefrontal Cortex in Ego Dissolution and Emotional Arousal During the Psychedelic State"

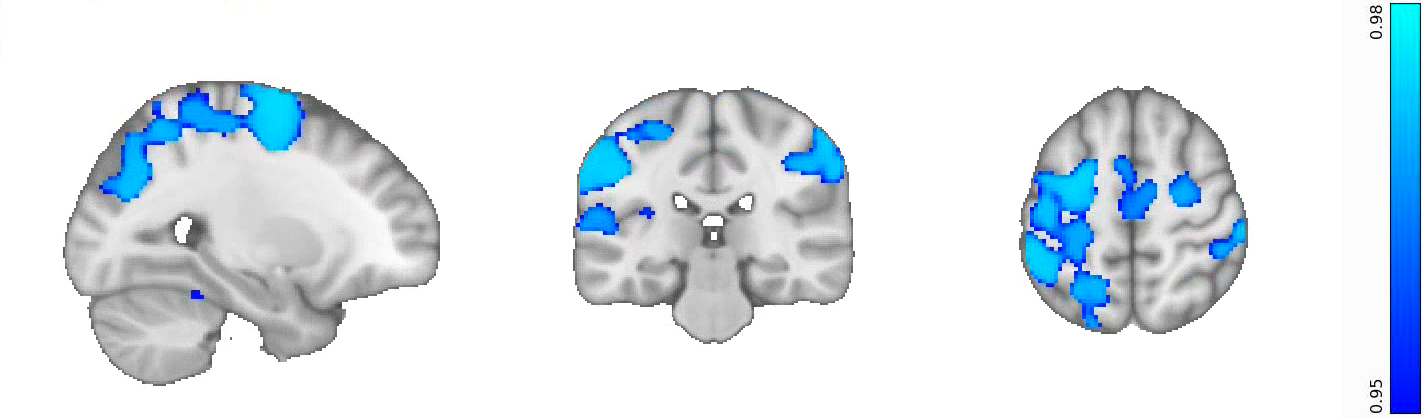

**Figure S1: Delta analysis reveals regions of the brain that become less functionally connected to the left DLPFC on LSD. Blue areas denote regions that decrease in RSFC on LSD; they include the left precentral gyrus, right supramarginal gyrus, left inferior frontal gyrus, and left fusiform gyrus.**

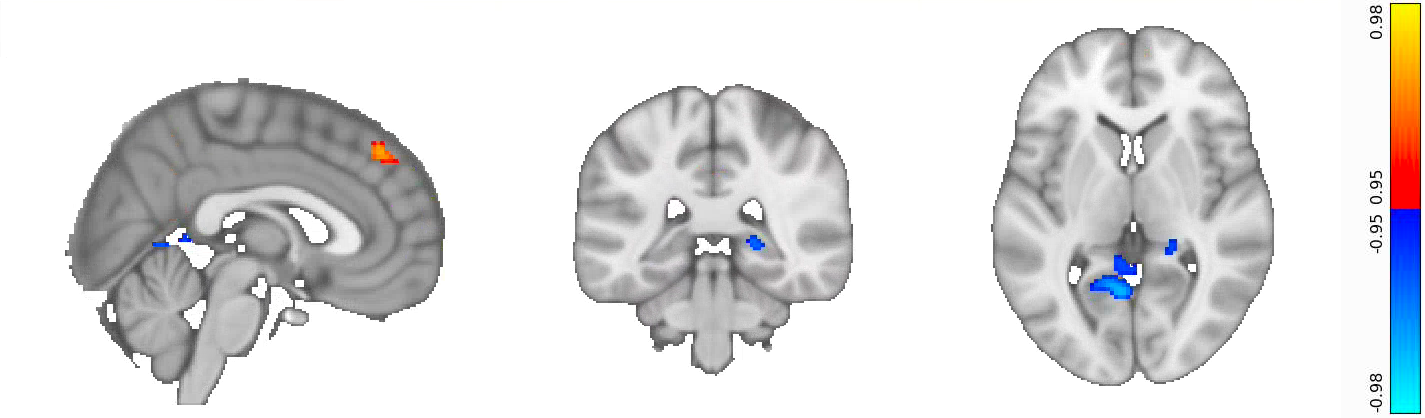

**Figure S2: Delta analysis reveals regions of the brain that become more or less functionally connected to the combined Thalamus and FFA seed from the ‘reverse’ ego dissolution analysis. Regions that increase in RSFC on LSD are shown in red/yellow; they include key hubs of the default mode network, i.e., the medial prefrontal cortex and posterior cingulate cortex. Blue areas denote regions that decrease in RSFC on LSD; they include the retrosplenial cortex, right parahippocampal gyrus, and right precuneus.**

Emotional Arousal

Demeand VAS Report

| RSFC Changes | Emotion Demean |
| --- | --- |
| -1.1437 | -7.4 |
| -0.0498244 | 1.1 |
| -2.02349 | -7.4 |
| 0.827651 | 0.1 |
| 0.574124 | 1.6 |
| -0.170683 | -4.9 |
| 1.06765 | 1.6 |
| -1.1577 | -0.9 |
| -1.06341 | -5.4 |
| 0.00451936 | -0.4 |
| -0.242934 | -0.9 |
| 1.31564 | 8.6 |
| -0.07098 | 4.1 |
| 1.37418 | 7.1 |
| 0.908595 | 3.1 |

Confirmational Emotional Arousal

| RSFC Changes | Demeand VAS Report |
| --- | --- |
| -0.39032 | -7.4 |
| 0.198557 | 1.1 |
| -1.42562 | -7.4 |
| 0.193998 | 0.1 |
| 0.187064 | 1.6 |
| -0.988581 | -4.9 |
| 0.189124 | 1.6 |
| -0.543994 | -0.9 |
| -1.574 | -5.4 |
| 0.00305288 | -0.4 |
| 0.093535 | -0.9 |
| 0.974331 | 8.6 |
| 0.195089 | 4.1 |
| 0.69094 | 7.1 |
| 0.380746 | 3.1 |

**S3 Figure: A scatterplot of the correlation between Emotional Arousal and the RSFC in the initial and confirmational analyses. The x-axis is the changes in RSFC between placebo and LSD. The y-axis is the demeaned VAS measures of Emotional Arousal taken post scan.**

Ego Dissolution

| RSFC Changes | Demaened VAS Report |
| --- | --- |
| 0.0382764 | -5.73 |
| 0.763111 | 10.27 |
| -0.223733 | -5.73 |
| -0.288149 | -3.73 |
| -0.112172 | -5.73 |
| -0.167511 | 0.27 |
| -0.416666 | -4.73 |
| -0.000851954 | -5.73 |
| -0.303437 | -5.73 |
| -0.626133 | -5.73 |
| -0.0634927 | -2.73 |
| 0.992121 | 14.27 |
| 0.0496522 | 4.77 |
| 0.513435 | 10.77 |
| 0.0254232 | 5.27 |

Confirmational Ego Dissolution

| RSFC Changes | Demeand VAS Report |
| --- | --- |
| -0.0822264 | -5.73 |
| 1.62403 | 10.27 |
| -0.871355 | -5.73 |
| -0.267193 | -3.73 |
| -0.0667869 | -5.73 |
| 0.322662 | 0.27 |
| -0.998587 | -4.73 |
| 0.0019012 | -5.73 |
| -0.845262 | -5.73 |
| -1.40471 | -5.73 |
| -0.389708 | -2.73 |
| 1.99126 | 14.27 |
| 0.277968 | 4.77 |
| 0.926364 | 10.77 |
| 0.864305 | 5.27 |

**S4 Figure: A scatterplot of the correlation between ego dissolution and the RSFC in the initial and confirmational analyses. The x-axis is the changes in RSFC between placebo and LSD. The y-axis is the demeaned VAS measures of Ego Dissolution taken post scan.**
